# Cofilin Drives Rapid Turnover and Fluidization of Entangled F-actin

**DOI:** 10.1101/156224

**Authors:** Patrick M. McCall, Frederick C. MacKintosh, David R. Kovar, Margaret L. Gardel

## Abstract

The shape of most animal cells is controlled by the actin cortex, a thin, isotropic network of dynamic actin filaments (F-actin) situated just beneath the plasma membrane. The cortex is held far from equilibrium by both active stresses and turnover: Myosin-II molecular motors drive deformations required for cell division, migration, and tissue morphogenesis, while turnover of the molecular components of the actin cortex relax stress and facilitate network reorganization. While many aspects of F-actin network viscoelasticity are well-characterized in the presence and absence of motor activity, a mechanistic understanding of how non-equilibrium actin turnover contributes to stress relaxation is still lacking. To address this, we developed a reconstituted in vitro system wherein the steady-state length and turnover rate of F-actin in entangled solutions are controlled by the actin regulatory proteins cofilin, profilin, and formin, which sever, recycle, and nucleate filaments, respectively. Cofilin-mediated severing accelerates the turnover and spatial reorganization of F-actin, without significant changes to filament length. Microrheology measurements demonstrate that cofilin-mediated severing is a single-timescale mode of stress relaxation that tunes the low-frequency viscosity over two orders of magnitude. These findings serve as the foundation for understanding the mechanics of more physiological F-actin networks with turnover, and inform an updated microscopic model of single-filament turnover. They also demonstrate that polymer activity, in the form of ATP hydrolysis on F-actin coupled to nucleotide-dependent cofilin binding, is sufficient to generate a form of active matter wherein asymmetric filament disassembly preserves filament number in spite of sustained severing.

**Significance Statement:** When an animal cell moves or divides, a disordered network of actin filaments (F-actin) plays a central role in controlling the resulting changes in cell shape. While it is known that continual turnover of F-actin by cofilin-mediated severing aids in reorganization of the cellular cytoskeleton, it is unclear how the turnover of structural elements alters the mechanical properties of the network. Here we show that severing of F-actin by cofilin results in a stress relaxation mechanism in entangled solutions characterized by a single-timescale set by the severing rate. Additionally, we identify ATP hydrolysis and nucleotide-dependent cofilin binding as sufficient ingredients to generate a non-equilibrium steady-state in which asymmetric F-actin disassembly preserves filament number in spite of sustained severing.

## Introduction

The capacity of the cytoskeletal protein actin to dynamically assemble into semi-flexible filaments (F-actin) underlies its fundamental role in cell motility (Pollard and Borisy, 2003), morphogenesis (Lecuit et al., 2011), division (Pollard, 2010), and mechanics (Blanchoin et al., 2014). More than 100 actin-binding proteins control the formation and differential regulation of F-actin networks, yielding architectures and turnover rates tuned for specific cellular processes (Pollard, 2016). Accordingly, the actin cortex, a thin and approximately isotropic F-actin meshwork anchored just beneath the plasma membrane is thought to be the primary determinant of cell shape and mechanics (Clark et al., 2013; Salbreux et al., 2012). In vivo measurements place cortical actin turnover on the 10-100 s timescale (Fritzsche et al., 2013; Robin et al., 2014), and while turnover is known to modulate cortical tension (Tinevez et al., 2009) and flows (Bray and White, 1988), a mechanistic understanding of how turnover regulates cortical mechanics is currently lacking.

Rheological measurements of F-actin networks reconstituted with purified proteins provide significant insight into the mechanics of living cells by enabling architectural and compositional control, and are sufficient to capture aspects of cellular mechanical response (Gardel et al., 2006). Guided by decades of reconstitution experiments, a quantitative theoretical understanding has emerged for how static microscopic parameters like filament density, length, and stiffness contribute to the viscoelastic mechanics of entangled F-actin solutions (Morse, 1998a, 1998b) and cross-linked F-actin networks (Broedersz and MacKintosh, 2014). Suppression of filament bending fluctuations by entanglements or cross-links transiently stores stress energy, giving rise to elasticity on the timescale of seconds. Diffusive, snake-like reptation of filaments (in entangled solutions) or cross-link unbinding (in networks) sets the timescale for stress relaxation, t_relax._ While the response of networks on timescales longer than t_relax_ is complicated by a broad spectrum of timescales related to the unbinding of multiple cross-links (Broedersz et al., 2010), relaxation is expected to be nearly Maxwellian in entangled solutions, with response dominated by a simple viscosity (Isambert and Maggs, 1996; Morse, 1998a). However, the contribution of dynamic F-actin turnover to stress relaxation remains largely unknown.

F-actin turnover requires sequential disassembly, nucleotide exchange, and assembly, and is limited *in vitro* primarily by slow disassembly kinetics (Brieher, 2013; Pollard, 1986). However, all of these reactions are tightly regulated in vivo, with the actin-binding proteins cofilin and profilin playing particularly important roles (Bugyi and Carlier, 2010; Pollard et al., 2000). Cofilin binds cooperatively (De La Cruz, 2005) and preferentially (Blanchoin and Pollard, 1999; Carlier et al., 1997) to ADP-F-actin, allosterically accelerates release of inorganic phosphate (Blanchoin and Pollard, 1999; Carlier et al., 1997; Suarez et al., 2011), and severs filaments at boundaries between clusters of cofilin-bound and -unbound subunits (De La Cruz, 2009; McCullough et al., 2011; Suarez et al., 2011). Profilin, in turn, competes with cofilin for binding to ADP-bound monomers (Blanchoin and Pollard, 1998), catalyzes exchange of ADP for ATP (Mockrin and Korn, 1980), blocks assembly at pointed-ends (Pollard and Cooper, 1984; Tilney et al., 1983), and promotes the rapid elongation of barbed-ends by formin proteins (Goode and Eck, 2007), which are responsible for generating the long cortical filaments important for mechanics in living cells (Fritzsche et al., 2016).

Here, we use purified cofilin, profilin, and formin to reconstitute rapid F-actin turnover at steady-state. We then combine filament-level measurements of length and turnover rate with fluorescence recovery after photobleaching (FRAP) and microrheology to systematically study the impact of non-equilibrium turnover on the dynamics and mechanics of entangled actin filament solutions *in vitro*. The choice of entangled solutions enables a quantitative assessment of stress relaxation mechanisms. Our work lays the foundation for elucidating the influence of F-actin turnover on the mechanics of more physiological network architectures.

## Results

### Independent control of F-actin length and turnover at steady-state in vitro

Nucleotide hydrolysis is intimately coupled to actin polymer dynamics, as shown in Fig. 1A. Upon incorporation into filaments, ATP-bound globular actin (ATP-G-actin) monomers undergo a conformational change, becoming ATP-F-actin. ATP is rapidly and stochastically hydrolyzed on the filament (Jégou et al., 2011; Korn et al., 1987), converting ATP-F-actin to ADP-Pi-F-actin (orange). The inorganic phosphate (Pi) is subsequently released on a much slower timescale (300-500 s) (Blanchoin and Pollard, 1999; Melki et al., 1996), resulting in ADP-F-actin (yellow). ADP-F-actin converts to ADP-G-actin upon dissociation from the filament, and the cycle is completed by exchange of the bound ADP nucleotide with free ATP in solution. Importantly, it is the effective irreversibility of the ATP-hydrolysis step which confers non-equilibrium dynamics to this set of coupled reactions, generating a directed turnover cycle with a steady-state flux of monomers. Monitoring the production of P_i_ in solution with a coupled-enzyme reaction gives a direct measure of bulk turnover (Webb, 1992).

**Figure 1.**
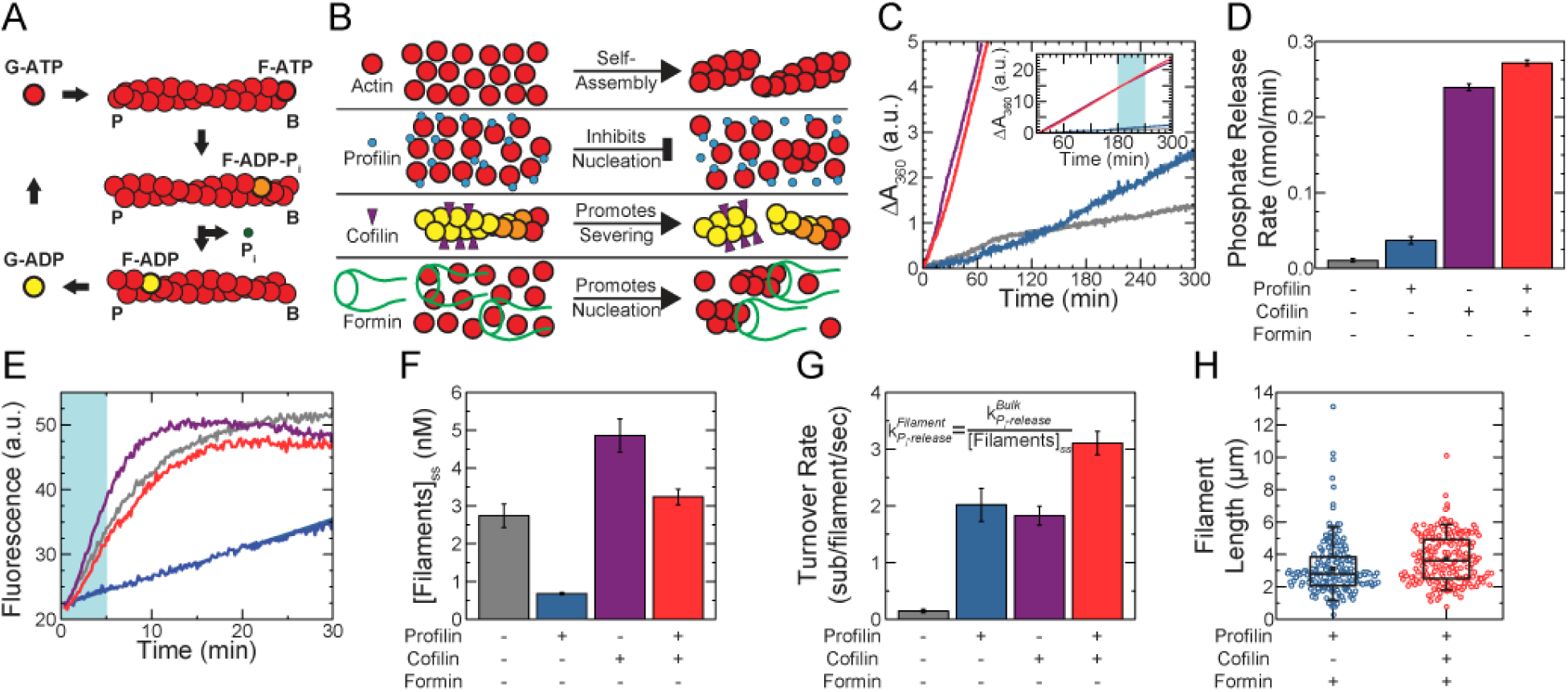
Independent control of actin length and turnover at steady-state *in vitro*. (A) Schematic of actin turnover cycle. B and P denote filament barbed and pointed ends, respectively. (B) Schematic of important biochemical activities of actin, profilin, cofilin, and formin. (C) Timecourse of inorganic phosphate (Pi) production for 1.5 μM Mg-ATP-actin alone (gray), with 4.5 μM profilin (R_p_ = 3, blue), with 0.75 μM profilin (R_c_ = 0.5, purple), or with 4.5 μM profilin and 0.75 μM cofilin (red), all in the presence of 0.2 mM MESG and 1 U/ml PNP. (Inset) Extended vertical axis showing the linear increase in all samples at long times. (D) Bulk phosphate release rate from linear fits to the Pi-release time courses during the time-window shaded in light blue (C, inset). Error bars indicate SEM, N = 4 for the profilin + cofilin condition, and N = 2 for each of the others. (E) Typical time courses of seeded assembly reactions in which 0.25 μM Mg-ATP-actin monomers (10 % pyrene-labeled) are added to 0.5 μM unlabeled actin seeds formed in the absence of additional proteins (gray), with 1.5 μM profilin (R_p_ = 3, blue), with 0.25 μM cofilin (R_c_ = 0.5, purple), or with 1.5 μM profilin and 0.25 μM cofilin (red). (F) Steady-state filament concentrations calculated using linear fits to the seeded assembly time courses during the time-window shaded in light blue in (E). Error bars indicate SEM, N = 3 for each condition. (G) Single filament turnover rates calculated from the data in (D,F) and rescaled by actin concentration for actin alone, with R_p_ = 3, with R_c_ = 0.5, or with R_p_ = 3 and R_c_ = 0.5. (H) Distribution of Alexa488-phalloidin-labeled filaments from source solutions containing 11.9 μM actin, R_p_ = 3, R_f_ = 0.01, and either no cofilin (R_c_ = 0, blue), or 6.95 μM cofilin (R_c_ = 0.5, red). Each length distribution is composed of 100 filaments from each of two independent samples, for a total of 200 filaments/condition.

F-actin turnover is regulated in part by the actin binding proteins profilin and cofilin (Fig. 1B) (Blanchoin and Pollard, 1999; Carlier et al., 1997; Didry et al., 1998). We measured turnover (inorganic phosphate (Pi) production; Fig. 1C) in solutions assembled from 1.5 μM Mg-ATP-actin alone (gray trace), or copolymerized with either 4.5 μM profilin (mole ratio profilin:actin = R_p_ = 3, blue), 0.75 μM cofilin (R_c_ = 0.5, purple), or 4.5 μM profilin and 0.75 μM cofilin (R_p_ = 3 and R_c_ = 0.5, red). All traces are initially non-linear as actin is assembled, but become linear once steady-state is reached (Fig. 1C, Fig. S1). Measurements of steady-state Pi production (180-240 min after initiation of polymerization) indicate that a molar excess of profilin:actin (R_p_ = 3) is sufficient to increase the bulk turnover rate ~3-fold over actin alone (Fig 1D). Optimal concentrations of cofilin (R_c_ = 0.5, determined from FRAP and microrheology, Figs 3-4) accelerate bulk turnover ~20-fold over actin alone, consistent with previous work (Carlier et al., 1997), while the combined effects of profilin and cofilin increase bulk turnover ~23-fold (red), qualitatively consistent with previous results (Didry et al., 1998).

To extract how profilin and cofilin affect the turnover rate of individual filaments, we first determined the number of filaments in solution from a variant on the seeded pyrene-actin assembly assay (Fig. 1E-F, SI Methods). Briefly, unlabeled actin is assembled in the presence of regulatory proteins and allowed to reach steady-state. Fluorescent pyrene-actin is then added, and, as in traditional seeded assays, the initial rate of the fluorescence increase is proportional to the number of elongating filaments present (Higgs et al., 1999).

Consistent with its role in inhibiting spontaneous nucleation (Pollard et al., 2000), profilin (R_p_ = 3) reduces the filament concentration at steady-state ~5-fold (Fig. 1B,E,F). By contrast, cofilin (R_c_ = 0.5) increases the filament concentration ~2-fold, qualitatively consistent with its severing activity (De La Cruz, 2009) (Fig. 1B,E,F). In the presence of both profilin and cofilin, the filament concentration is comparable to that for actin alone. By dividing the bulk phosphate release rate by the filament density, we obtain the turnover rate for individual filaments (Fig. 1G). We find that profilin and cofilin are each sufficient to increase the turnover rate of actin ~15-fold, from 0.14 to ~2.0 subunits/filament/sec. In the presence of both profilin and cofilin, turnover increases further to ~3.1 subunits/filament/sec, qualitatively consistent with previous results (Didry et al., 1998). Taken together, these data demonstrate that the combination of profilin and cofilin increases the turnover rate of individual actin filaments ~22-fold.

Filament length is controlled by varying the filament nucleation rate through changes in the concentration of the formin mDia1, which nucleates (Li and Higgs, 2003) and processively elongates (Kovar et al., 2006; Romero et al., 2004) actin filaments (Fig. 1B). Fluorescence imaging revealed that, for fixed concentrations of actin and profilin, the mean filament length at steady-state can be varied from 21 to 3 μm by increasing the formin concentration from 0 to ~1 μM (Fig S2). Remarkably, the presence of the severing protein cofilin has little affect on the steady-state length distribution in the presence of profilin and formin (Fig. 1H). Indeed, 11.9 μM Mg-ATP-actin polymerized in the presence of profilin (R_p_ = 3, 35.7 μM) and formin (R_f_ = 0.01, 119 nM) without (R_c_ = 0, blue) or with cofilin (R_c_ = 0.5, 5.95 μM, red) have very similar length distributions with mean lengths of 3.1 μm without cofilin and 3.7 μm in the presence of cofilin. Together these data demonstrate that nucleation during assembly sets the steady-state F-actin length distribution nearly independent of cofilin-mediated severing and increased filament turnover dynamics.

### Cofilin enhances reorganization dynamics in entangled actin solutions

To explore the consequences of severing and turnover on the dynamic redistribution of F-actin, we performed steady-state Fluorescence Recovery After Photobleaching (FRAP) measurements on entangled solutions of 11.9 μM actin (5% Oregon Green-actin) assembled with a constant molar excess of profilin (R_p_ = 3) and a range of cofilin and formin concentrations. At this actin concentration, the expected distance between actin filaments is ~ 420 nm, yielding spatially uniform actin fluorescence prior to bleaching (Fig. 2A).

**Figure 2.**
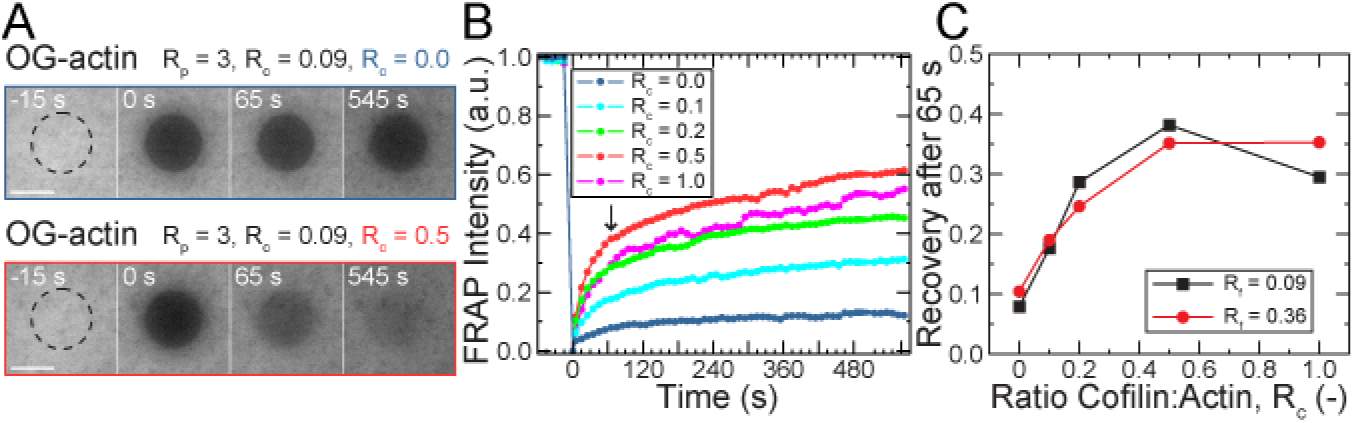
Cofilin enhances reorganization in entangled actin solutions. (A-C) All samples are polymerized from 11.9 μM Mg-ATP-actin (5 % Oregon Green-labeled) with R_p_ = 3 (35.7 μM), R_f_ = 0.09 (1.07 μM), and R_c_ as indicated (0-11.9 μM), except for the red circles in (C), for which R_f_ = 0.36 (4.28 μM). (A) Confocal fluorescence time-lapse micrographs with R_c_ = 0 (B, blue) or R_c_ = 0.5 (C, red) at steady-state. A dashed black circle denotes the region exposed to the bleaching laser. Time is relative to the end of the 5-s bleach. Scale bar is 50 μm. (B) Time course of the normalized fluorescence intensity averaged across the bleached region for entangled solutions with R_c_ as indicated in the legend and R_f_ = 0.09. (C) Actin fluorescence intensity recovered within 65 s of bleaching (denoted by arrow in (B)) as a function of cofilin concentration for two concentrations of formin. Each point represents a single experiment.

The post-bleach recovery of actin fluorescence we observe is qualitatively described by two phases: a relatively rapid recovery in the first ~60 s, and a much slower recovery thereafter (Fig. 2B). In the absence of cofilin, the average fluorescence intensity of the bleached region recovers to approximately 8 % of the pre-bleach value within 65 seconds, and then only to 12 % after 545 seconds (blue). By contrast, the recovery is much more rapid in the presence of cofilin (Fig. 2A-B, red), reaching approximately 40 % in 65 seconds and more than 60 % by 545 seconds for R_c_ = 0.5. Changes in the diffusivity of monomers or filaments are insufficient to fully account for this recovery profile, since the presence of cofilin increases the size of the monomer pool from ~1 % to only ~9 % total actin (SI Text), and cofilin doesn’t reduce the mean filament length (Fig. 1H). Instead, we interpret the pronounced recovery enhancement at early times as resulting primarily from accelerated F-actin turnover (Fig. 1G), where the rates at which unbleached subunits are incorporated and bleached subunits are removed from existing filaments are elevated through the action of cofilin.

The degree of actin fluorescence recovery 65-seconds after bleaching is sensitively tuned by cofilin (Fig. 2C), with a maximal increase of ~5-fold. Interestingly, on this timescale, we find comparable cofilin-dependent recovery for samples with different concentrations of formin (black squares and red circle). Since the mean length is expected to differ by a factor of 2 between these formin concentrations (SI Text), the near independence of the fluorescence recovery to formin concentration supports that notion that filament diffusion plays a minimal role in the recovery on this timescale. The slow diffusion of long filaments is likely responsible for the incomplete recovery after 545 s observed in all cases, however. Taken together, these data demonstrate that *in vitro* F-actin turnover, catalyzed by cofilin and relying on monomer diffusion, is the dominant process controlling the steady-state reorganization of F-actin on the timescale of 10s of seconds.

### Cofilin fluidizes entangled F-actin solutions

We employ microrheology to measure the frequency-dependent viscoelasticity of entangled F-actin solutions with varying concentrations of cofilin at steady-state. 11.9 μM Mg-ATP-Actin is assembled with the desired concentrations of profilin, formin, cofilin, and 1-μm diameter fluorescent polystyrene beads for 95 minutes to reach steady-state (Fig. S4). From fluorescent images, the bead centroids are tracked over time and the ensemble-averaged mean-squared-displacements (MSDs) as a function of lag time (Δt) are calculated. In the presence of saturating profilin (R_p_ = 3) and moderate formin (R_f_ = 0.1), but the absence of cofilin, bead MSDs are characteristic of that observed in semi-dilute, entangled F-actin solutions (Fig. 3A, blue). For times less than 0.3 s, the time-dependence of the MSD arises from bending fluctuations of individual F-actin (Amblard et al., 1996; Broedersz and MacKintosh, 2014). At longer times, the MSD approaches a plateau value that reflects the local elastic modulus (Gardel et al., 2003). The local viscoelasticity can be measured by using the Generalized Stokes-Einstein Relationship to obtain the frequency-dependent elastic, G’, and viscous, G”, moduli (Gardel et al., 2003; Gittes et al., 1997; Squires and Mason, 2010). The elastic modulus is nearly constant and much larger than the viscous modulus at frequencies between ~1 Hz and 0.01 Hz. This is indicative of a material that is predominantly elastic over this frequency regime, consistent with previous measurements on entangled F-actin solutions (Gardel et al., 2003; Liu et al., 2006).

**Figure 3.**
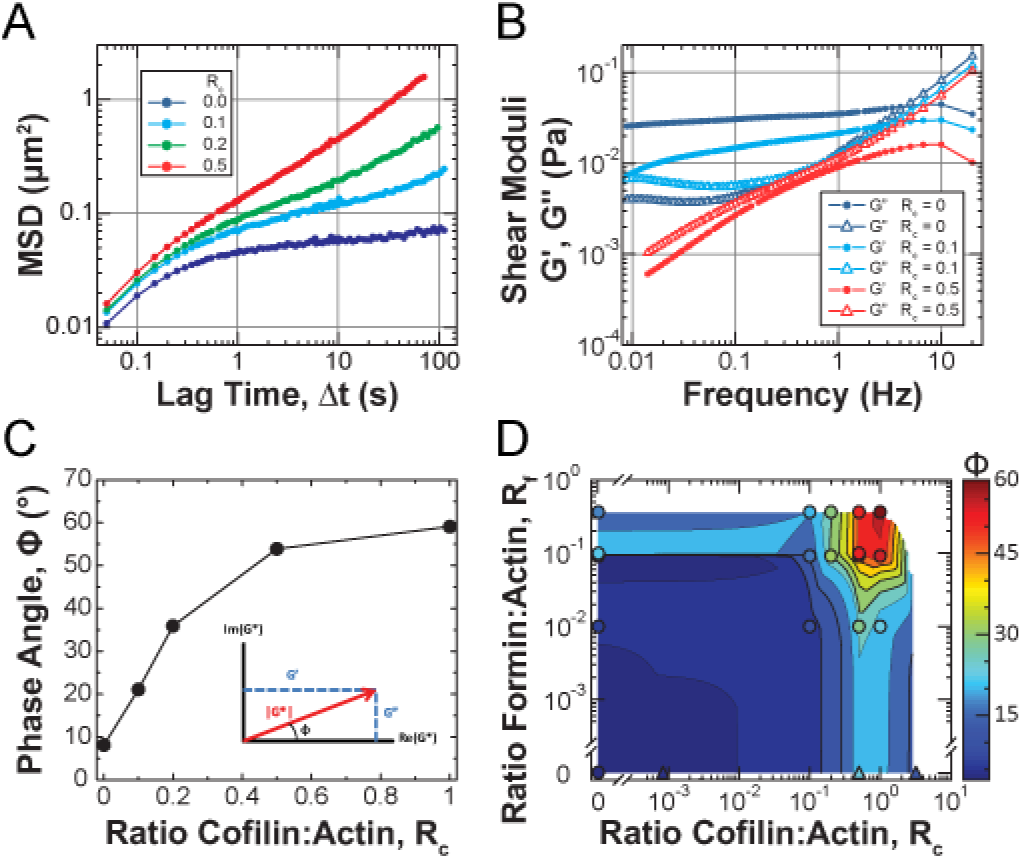
Cofilin-mediated turnover tunes the steady-state fluidity of entangled F-actin solutions. (A-D) All microrheology measurements are of steady-state entangled actin solutions polymerized from 11.9 μM Mg-ATP-actin (0 % or 5 % Oregon-Green labeled) with R_p_ = 3 (35.7 μM), R_f_ = 0.09 (1.07 μM), and R_c_ as indicated, except for (D), where R_f_ is as indicated, and R_p_ = 0 for samples denoted by triangles. (A) Ensemble-averaged mean-squared-displacement (MSD) of 1-um diameter beads with R_c_ as indicated in the legend. Each point is calculated from at least 1000 displacements from a single sample. (B) Real and imaginary components of the complex shear modulus (G’ and G”, respectively) for the R_c_ = 0, 0.1, and 0.5 samples from (A), denoted by closed circles (G’) and open triangle (G”), respectively. (C) Phase angle (Φ) evaluated at 0.1 Hz for conditions with R_c_ as indicated. (Inset) Geometric relationship between the magnitude of the complex shear modulus (|G*|, red), its real and imaginary components (G’, G”, blue) and the phase angle (Φ), shown in the complex plane. (D) State diagram displaying the phase angle (Φ, color) evaluated at 0.1 Hz for entangled solutions with R_f_ and R_c_ as indicated. In most cases, each point represents a single sample, though some are the average of multiple samples. The heatmap represents a 2D interpolation of the discrete data points.

Increasing concentrations of cofilin raises the magnitude and modifies the time-dependence of the MSD. The plateau in the MSD is truncated by the emergence of a gradual upturn at lag times greater than 10 s for R_c_ = 0.1 (Fig 3A, cyan). The location of the upturn shifts toward shorter lag times as R_c_ increases, with the MSD approaching diffusive scaling (~ *t^1.0^*) at the longest lag times for R_c_ = 0.5. This increased bead mobility reflects dramatic changes in the local viscoelasticity (Fig. 3B). At moderate cofilin concentration (R_c_ = 0.1, cyan), the elastic modulus systematically decays from 1 to 0.01 Hz, resulting in a low frequency crossover where, presumably, the viscous modulus becomes dominant at frequencies below 0.1 Hz. At higher cofilin concentrations (R_c_ = 0.5), the elastic and viscous modulus are similar in magnitude and decay with time. A parameterization of the frequency-dependent viscoelasticity is the phase angle Φ, or arc tangent of the ratio G”/G’, and would be 0° and 90° for purely elastic and viscous materials, respectively, at a given frequency. Calculating Φ at 0.1 Hz, we find that it increases from 10° to 60° as the cofilin concentration is increased from R_c_ = 0 to R_c_ = 1 (Fig. 3C). Thus, increased cofilin concentration results in a transition between a viscoelastic solid to a viscoelastic fluid. Since cofilin is not reducing the mean filament length (Fig 1H), and since the steady-state monomer pool is not increased sufficiently to increase the mesh size beyond the radius of the bead (SI text), the fluidization likely results from elevated actin filament turnover.

To explore how the fluidity of actin solutions can be regulated by changes to filament length and turnover dynamics, we measure Φ at 0.1 Hz over a range of cofilin and formin concentrations (Fig. 3D). For all filament lengths (formin concentrations) examined, the addition of sufficient cofilin increases the phase angle. The most fluid-like samples are those with short filaments (highest formin concentration) undergoing rapid turnover (high cofilin concentration). Interestingly, for the longest filaments, we see a hint that the phase angle a biphasic function of cofilin concentration, peaking near R_c_ = 0.5-1, and decreasing above R_c_ ~1, implicating the severing activity of cofilin (De La Cruz, 2009; McCullough et al., 2011; Suarez et al., 2011) in driving the fluidization. While fluidization of entangled F-actin solutions by shortening the steady-state filament length has been previously appreciated (Gardel et al., 2003), we demonstrate here that by accelerating steady-state turnover, fluidization can also be achieved without reducing the average length of filaments.

### Rapid cofilin-mediated turnover is a single-timescale mode of stress relaxation

To compare the mechanism of enhanced fluidity that arises from accelerated filament turnover to shortened filament length, we compare the MSD of microscopically distinct F-actin solutions that have identical values of Φ at 0.1 Hz (Fig. 4A). Specifically, we compare samples with long filaments and fast turnover (purple, red) to one with short filaments and slow turnover (black). Interestingly, these samples are rheologically indistinguishable at all timescales probed. Thus, an entangled solution of relatively long filaments undergoing rapid turnover (purple) is mechanically equivalent to a solution of relatively short, stable filaments (black).

**Figure 4.**
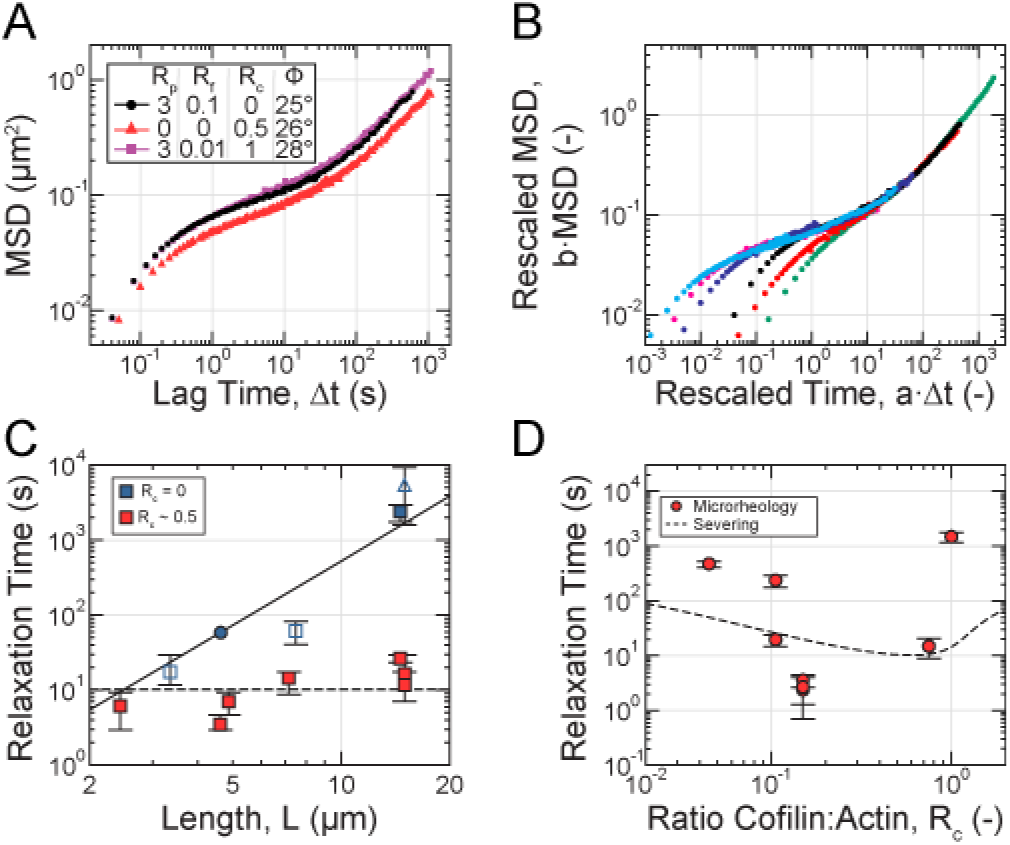
Rapid cofilin-mediated turnover is a single-timescale mode of stress relaxation and dominates reptation. (A-D) All microrheology measurements are of steady-state entangled actin solutions polymerized from 11.9 μM Mg-ATP-actin (0 % or 5 % Oregon-Green labeled) with R_p_ = 3 (35.7 μM), R_f_ and R_c_ as indicated, except for the sample denoted by triangles in (A,C), where R_p_ = 0. (A) Ensemble-averaged MSD for samples with similar values of the phase angle (Φ) evaluated at 0.1 Hz. (B) Collapse of MSD curves at long times to a reference sample of short stable filaments (A-B, black; C, circle) after rescaling lag time and MSD by shift factors *a* and *b*, respectively. (C) Dependence of relaxation time, estimated from the shift factor *a*, on filament length with (red) and without (blue) cofilin. Closed and open symbols represent measurements from a single sample and from two or more samples, respectively. Solid curve is the relaxation time predicted for reptation alone. Dashed line is a fit of the length-independent severing-based model to the cofilin-containing data. (D) Dependence of relaxation time on R_c_. Dashed line is the relaxation time dependence on cofilin concentration predicted by the severing-based model, using the severing rate inferred from the fit in (C), association constant for cofilin:F-actin binding of K_a_ = 1/10 μM, and a cofilin-binding cooperativity parameter of ω = 7.5 (De La Cruz, 2005).

Notably, such global equivalence in the mechanical response requires equivalence in the long-timescale relaxation dynamics, despite the differences in underlying microscopic processes. Since the stress relaxation of entangled solutions is dominated by a single timescale (Morse, 1998b), we infer that the enhanced relaxation resulting from cofilin-mediated actin turnover is also a single-timescale process. In a manner analogous to time-temperature superposition (Ferry, 1980; Tobolsky, 1956), we estimate the relaxation timescale for individual F-actin solutions by rescaling the measured MSDs by a shift-factor *b*, and the separation time Δt by a shift-factor *a*, such that the long-time behavior superposes with that of a single reference sample composed of relatively short filaments in the absence of cofilin (black, Fig. 4A-B), for which the relaxation time, *τ_ref_*, could be directly inferred from the low-frequency crossover of G’ and G” (Fig. S5). The estimated relaxation timescale for each sample *i* is then given by *τ_i_* = *τ_ref_*/*α_i_*. The successful superposition of the long-timescale MSDs for entangled solutions with a wide variety of filament lengths and cofilin concentrations (Fig. 4B) validates the use of this approach.

We examine the dependence of the relaxation time on filament length, which is controlled by tuning the formin concentration. Absolute values of the mean filament length are inferred from a mathematical model which incorporates formin concentration and nucleation rate constants extracted from spontaneous assembly measurements (SI Text). In the absence of cofilin, the increase in relaxation time as a function of *L* is consistent with *L*^3^ (Fig. 4C, blue). This scaling is that predicted for stress relaxation via reptation (Doi and Edwards, 1986), roughly the time it takes a filament to diffuse its own length. This is the mechanism of stress-relaxation in entangled actin solutions (Isambert and Maggs, 1996; Morse, 1998a, 1998b). By contrast, at optimal cofilin concentrations, the relaxation time is reduced at all values of filament lengths examined, and exhibits much weaker scaling, consistent with linear, or even sub-linear, dependence on filament length (Fig. 4C, red). This indicates that cofilin-mediated turnover accelerates stress relaxation in entangled solutions by a mechanism distinct from reptation. Taken together, these data demonstrate that cofilin-meditated turnover results in a single-timescale mode of stress relaxation that can dominate over filament reptation.

We explore how changing the severing rate can modulate the stress relaxation time. As the cofilin molar ratio is increased from 0.05 to 0.15 (towards the optimal concentration), we see that the relaxation time decreases nearly ~200-fold from 600 s to ~3 s (Fig 4D). As the cofilin ratio is increased further to 1, the relaxation time increases back to ~1000 s. This biphasic dependence of stress relaxation time on cofilin concentration is qualitatively consistent with the biphasic dependence of severing rate on cofilin binding density (Andrianantoandro and Pollard, 2006; De La Cruz, 2005, 2009; Suarez et al., 2011) (Fig 4D, dashed line). Importantly, this data underscores how non-equilibrium severing activity can decouple mechanical stress relaxation from material structure, as all of these samples contain actin filaments at the same density and nearly the same length.

To understand the mechanism of cofilin-mediated stress relaxation observed in Fig. 4C and D, we explore a simple physical model which explicitly incorporates filament severing, and which captures the weak dependence of the relaxation time on filament length. Assuming an initial steady-state length distribution 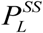, we approximate the residual stress at time *t* following application of a step strain as proportional to the residual polymer of length exceeding the entanglement length 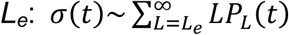. The distribution *P_L_*(*t*) of residual length evolves according to 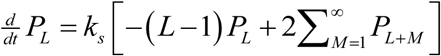 due to severing alone at a rate *k_s_* and neglecting depolymerization, which represents a valid approximation for large lengths *L* and *M*. This time evolution is analogous to that used previously to describe the rheology of worm-like micelles (Cates, 1987), but neglecting filament annealing reactions, which should be suppressed here by depolymerization and formins (Kovar et al., 2003). We emphasize that *P_L_*(*t*) represents the distribution of “stressed” filaments, and not the instantaneous filament length distribution 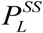, which differs due to depolymerization that leads to stress-free elongation. This model is exactly solvable for exponential 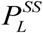 (SI Text), and interestingly yields a stress that decays with a characteristic timescale *τ_relax_* − (*k_s_L_e_*)^−1^ which depends on polymer density (through the entanglement length) but not filament length (Fig S6).

A fit of this length-independent model to the relaxation times measured with cofilin (Fig. 4C, dashed line) predicts a severing rate of ~3.3 x 10^-4^ events/sub/s at optimal concentrations. Although ~12-fold higher than rates measured for isolated, unconstrained filaments (McCullough et al., 2011), constraints like those introduced by entanglements are known to enhance severing by cofilin (De La Cruz et al., 2015; Pavlov et al., 2007). Including depolymerization in the evolution of *PL* could further reduce this discrepancy, and likely contributes to the differences in Fig. 4D. Interestingly, one can define a lengthscale 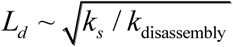 at which the time between severing events is comparable to the time for depolymerization of fragments of length *L_d_*. If this length exceeds the entanglement length, then the rate of stress relaxation is expected to increase by a factor of ~*L_d_*/*L_e_*. Elucidation of the role of this lengthscale in the dynamics of stress relaxation is currently under way. Taken together, however, these modeling results support severing as a crucial ingredient for the weak dependence of mechanical relaxation on filament length.

## Discussion

Here we show that cofilin-mediated turnover tunes the steady-state fluidity of entangled solutions of F-actin. In the presence of formin and profilin, cofilin-mediated severing does not reduce the mean filament length or number density at steady-state, factors known to control mechanics and filament mobility. Instead, the enhanced fluidity arises from non-equilibrium actin turnover catalyzed by cofilin severing.

Our work provides a new microscopic model of how a steady-state length is achieved in the presence of cofilin activity (Fig. 5). The textbook view of actin turnover is treadmilling, wherein barbed-end elongation proceeds at the turnover rate, and is exactly balanced at steady-state by pointed-end disassembly (Phillips et al., 2008). A treadmilling-based mechanism predicts that the relaxation time would be linear in L, which is the strongest dependence with which our data is reasonably consistent (Fig. 4D). However, the 8 nm/s treadmilling velocity we obtain from turnover measurements is too slow by more than 100-fold to account for the relaxation times we estimate (Fig. 4D).

**Figure 5.**
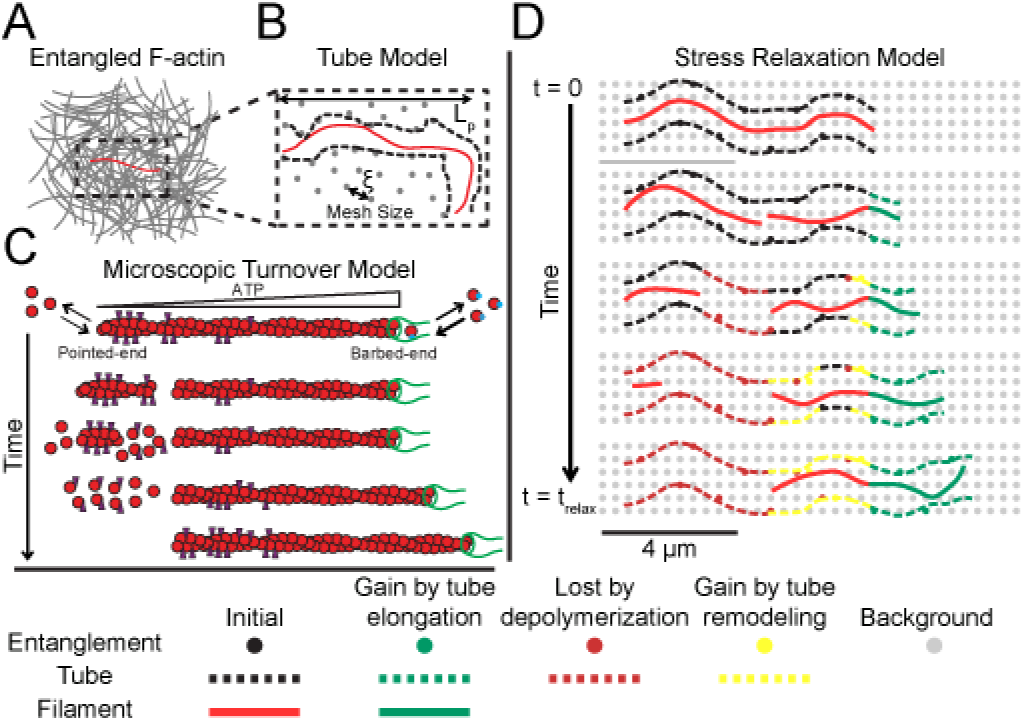
Microscopic model of cofilin-mediated actin turnover and stress relaxation. (A) Cartoon of a test filament (red) entangled with other filaments (gray) in a semi-dilute actin solution. (B) Representation of tube model. Entanglements (black) between the test chain and background filaments constrain test chain fluctuations to a tube (black dashed line) and thereby store stress. The entanglement length is set by solution mesh size (gray dots) and filament persistence length. (C) Microscopic dynamics of the test filament regulated by cofilin, profilin, and formin. Cofilin binds and severs ADP-rich filament regions, creating unstable ADP-rich filaments which depolymerize from both ends. The additional G-actin is incorporated onto the stable ATP-barbed end, restoring the length lost from severing. (D) Tube memory, and thus stress, decays by depolymerization (maroon) and tube remodeling (yellow) after severing of the test filament and background filaments, respectively. Polymerization (green) is stress-free, so newly created tube does not contribute.

Instead, non-equilibrium ATP hydrolysis directs cofilin severing and enables asymmetric kinetics for both assembly and disassembly of severed fragments (Fig. 5C). Cofilin binds with an ~40-fold higher affinity to ADP-F-actin than either ADP-Pi- or ATP-actin (Blanchoin and Pollard, 1998; Carlier et al., 1997), preferentially localizing both binding and severing away from the filament barbed (McCullough et al., 2011; Suarez et al., 2011). Importantly, the two fragments formed upon filament severing differ in their nucleotide composition. The fragment with the newly-created barbed end is ADP-rich throughout, resulting in its rapid disassembly from both pointed and barbed ends, at a combined rate on the order of ~0.1 μm/s (SI Text). In contrast, the fragment with the pre-existing barbed-end retains an ATP gradient along its length, and is thus stable, independent of the presence of the formin. This stable fragment continues to elongate, and by consuming monomer released through the rapid disassembly of unstable severed fragments, it quickly regenerates length lost through severing (Fig. 5C). Thus, severing couples to asymmetric (dis)assembly dynamics generated by non-equilibrium ATP-hydrolysis to preserve steady-state filament length while catalyzing turnover.

The possibility for non-equilibrium effects to fluidize materials is of great interest in understanding active biological materials. The mechanical response of entangled F-actin solutions is typically dominated by filament length and density, and understood in terms of a tube model (Fig 5A-B), wherein stress relaxes by filament reptation. Our results demonstrate that rapid turnover catalyzed by cofilin can fluidize these solutions without major changes to global architecture. The enhanced stress relaxation arises from rapid disassembly of large filament portions, while filament assembly and tube remodeling occur stress-free (Fig. 5D). In contrast to other active processes (e.g. myosin motors), here the non-equilibrium activity primarily effects the dynamical properties of the filament steady-state, enabling rapid (dis)assembly with no nucleation and ultimately enhancing stress relaxation, rather than altering local mechanics via generating local forces per say. This suggests that the bead motions are likely still thermally driven, implying that the fluctuation-dissipation theorem (Squires and Mason, 2010) is only weakly broken by the non-equilibrium turnover, and validating our use of the Generalized Stokes-Einstein Relation (Fig. 3).

It will be interesting to explore the effects of severing-mediated stress relaxation in more physiological cross-linked networks of F-actin. While saturation of actin networks with cross-linkers (Schmoller et al., 2011) or the side-binding protein tropomyosin (Christensen et al., 2017) inhibits disassembly in vitro, consistent with suppression of cofilin binding (De La Cruz, 2009), more sparsely distributed attachment points give both sufficient space to allow cofilin binding, and actually accelerate severing (Pavlov et al., 2007), suggesting an important role for crosslink density in tuning severing and thereby fluidity.

Finally, in cells, F-actin turnover has been considered an important mechanism to support the fluidization of the actin cortex on the scale of 1-60 s (Salbreux et al., 2012), but is mechanistically best understood in the context of protrusive structures like lamellipodia, which facilitate cell motility (Blanchoin et al., 2014). Turnover of polarized lamellipodial networks and listeria comet tails (Loisel et al., 1999; Theriot and Mitchison, 1991) is thought to proceed via a treadmilling array model (Pollard and Borisy, 2003) in which new filaments are continually nucleated near the leading edge, and elongate only briefly before being capped. Following transport away from the membrane by retrograde flow, capped filaments are then disassembled by cofilin and recycled by profilin, giving rise to spatially separated zones of assembly and disassembly. In contrast, a direct consequence of the dynamical regime of actin turnover we describe in the present work, which does not require steady-state filament nucleation, is that assembly and disassembly are spatially uniform. This mechanism may therefore be better-suited for isotropic networks like the cortex compared to polarized ones like the lamellipodia.

Future work exploring potential coupling between cofilin-mediated non-equilibrium turnover and network architecture, both in cells and reconstituted systems, will help elucidate how cells differentially tune the dynamics and mechanics of actin networks to facilitate distinct functions.

## Materials and Methods

Mg-ATP-actin (5% OG-labeled) was polymerized for 95 min in the presence of regulatory proteins and 1-μm diameter polystyrene beads to reach steady-state, and imaged on a spinning-disk confocal microscope. Details of all experimental methods and analysis can be found in the *SI Materials and Methods.*

## Acknowledgements

We are grateful to members of the M.L.G. and D.R.K. Laboratories, especially C. Suarez, D. Zimmermann, J. Winkelman, and P. Oakes, as well as W. McFadden, E. Munro, and E. De La Cruz for helpful discussions and suggestions. This work was supported by University of Chicago Materials Research Science and Engineering Center (NSF DMR-1420709).

